# Hypocretin Receptor 1 Blockade Early in Abstinence Prevents Incubation of Cocaine Seeking and Normalizes Dopamine Transmission

**DOI:** 10.1101/2024.11.30.625912

**Authors:** Philip J. Clark, Volodar M. Migovich, Sanjay Das, Wei Xi, Sandhya Kortagere, Rodrigo A. España

**Affiliations:** Department of Neurobiology and Anatomy, Drexel University College of Medicine, Philadelphia, PA 19129; Department of Microbiology and Immunology, Drexel University College of Medicine, Philadelphia, PA 19129

## Abstract

Abstinence from cocaine use has been shown to elicit a progressive intensification or incubation of cocaine craving/seeking that is posited to contribute to propensity for relapse. While the mechanisms underlying incubation of cocaine seeking remain elusive, considerable evidence suggests that abstinence from cocaine promotes mesolimbic dopamine adaptations that contribute to exaggerated cocaine seeking. Consequently, preventing these dopamine adaptations may reduce incubation of cocaine seeking and thereby reduce the likelihood of relapse. In the present studies, we first examined if incubation of cocaine seeking was associated with aberrant dopamine transmission in the nucleus accumbens after seven days of abstinence from intermittent access to cocaine. Given the extensive evidence that hypocretins/orexins regulate motivation for cocaine, we then examined to what extent hypocretin receptor 1 antagonism on the first day of abstinence prevented incubation of cocaine seeking and dopamine adaptations later in abstinence. Results indicated that abstinence from intermittent access to cocaine engendered robust incubation of cocaine seeking in both female and male rats. We also observed aberrant dopamine transmission, but only in rats that displayed incubation of cocaine seeking. Further, we showed that a single injection of the hypocretin receptor 1 antagonist, RTIOX-276, on the first day of abstinence prevented incubation of cocaine seeking and aberrant dopamine transmission. These findings suggest that hypocretin receptor 1 antagonism may serve as a viable therapeutic for reducing cocaine craving/seeking, thus reducing the likelihood of relapse.

## Introduction

Relapse to drug use after periods of abstinence remains a major obstacle in the treatment of substance use disorders. In humans, abstinence from drug use is associated with a progressive intensification or incubation of drug craving/seeking (Bedi et al., 2011; Wang et al., 2013; Li et al., 2015), which is hypothesized to promote relapse (Gawin and Kleber, 1986). This intensification of drug craving has been modeled preclinically as increases in cue-induced drug seeking across abstinence, leading to the hypothesis that abstinence may induce neuroadaptations that underlie the incubation of drug seeking (Grimm et al., 2001; Grimm et al., 2003; Bienkowski et al., 2004; Mead et al., 2007; Abdolahi et al., 2010) (Loweth et al., 2014; Terrier et al., 2016).

Accumulating evidence implicates the mesolimbic dopamine pathway in the development of incubation of cocaine seeking (Mameli et al., 2009; Sun and Wolf, 2009; Calipari et al., 2015). For example, recent work from our laboratory revealed that 28 days of abstinence following intermittent access (IntA) to cocaine promoted robust incubation of cocaine seeking and enhanced dopamine transporter (DAT) function in the NAc core (Alonso et al., 2022). These and related observations suggest that incubation of cocaine seeking may be linked to DAT adaptations and that normalizing dopamine transmission during periods of abstinence may be an effective strategy for reducing cocaine seeking. Unfortunately, pharmacotherapies that target dopamine systems directly, are largely ineffective or intolerable and may have abuse potential themselves (Minozzi et al., 2015).

Extensive evidence indicates that the hypocretin/orexin (Hcrt) peptides participate in reward and reinforcement and regulate dopamine transmission. For example, Hcrt-1 peptide increases motivation for cocaine (España et al., 2011) and reinstates cocaine-seeking (Boutrel et al., 2005; Matzeu et al., 2016), while Hcrt receptor 1 (Hcrtr1) antagonism decreases motivation for cocaine (Borgland et al., 2009; España et al., 2010; Muschamp et al., 2014; Bentzley and Aston-Jones, 2015; Brodnik et al., 2015; Prince et al., 2015; Levy et al., 2017), blocks reinstatement of cocaine-seeking (Aston-Jones et al., 2009; Smith et al., 2009; Mahler et al., 2013), and decreases behavioral sensitization to cocaine (Borgland et al., 2006). Moreover, knockout of Hcrt peptides reduces cocaine conditioned place preference (Shaw et al., 2017), and knockdown of Hcrtr1 reduces motivation for cocaine (Bernstein et al., 2017; Black et al., 2023).

The behavioral effects of Hcrt manipulations involve alterations in mesolimbic dopamine transmission. Hcrt neurons innervate the dopamine-rich ventral tegmental area (VTA), where Hcrt-1 potentiates excitatory drive and increases dopamine neuron firing (Peyron et al., 1998; Marcus et al., 2001; Fadel and Deutch, 2002; Korotkova et al., 2003; Borgland et al., 2006; Narita et al., 2006; Muschamp et al., 2007; Borgland et al., 2009; Muschamp et al., 2014). Further, acute administration of HcrtR1 antagonists decreases dopamine in the NAc under baseline conditions and reduces dopamine responses to cocaine (España et al., 2010; Prince et al., 2015; Levy et al., 2017; Brodnik et al., 2020a). Although these observations have been instrumental in establishing the involvement of Hcrt in drug-related reinforcement/motivation, to date there have been no reports examining the effects of Hcrtr1 disruption during periods of abstinence on incubation of drug seeking.

In these studies, we first examined to what extent IntA to cocaine promoted incubation of cocaine seeking and dopamine adaptations in the NAc after 7 days of abstinence. Female and male Long Evens rats self-administered cocaine on an IntA schedule and then underwent 7 days of abstinence. On abstinence day 1 (AD1) and abstinence day 8 (AD8), rats underwent cue-induced cocaine seeking tests to assess incubation of cocaine seeking. The following day, we used fast-scan cyclic voltammetry (FSCV) to examine if abstinence from IntA to cocaine promoted alterations in dopamine transmission and DAT expression and phosphorylation. In a separate group of rats, we explored the utility of a single injection of the Hcrtr1 antagonist, RTIOX-276, in preventing the incubation of cocaine seeking and DA adaptations observed after IntA to cocaine. Rats self-administered on the IntA schedule for 7 days and on AD1 were tested for cue-induced cocaine seeking. Immediately after the AD1 seeking test, rats received a single injection of vehicle or 20 mg/kg RTIOX-276 and were then left undisturbed. On AD8 rats were again tested for cue-induced seeking and the day following we performed FSCV and biochemistry to measure changes in dopamine transmission and DAT expression.

## Methods

### Animals

Adult female (220-260g) and male (300-410g) Long Evans rats (Envigo, Frederick, MD, USA) were maintained on a 12 h reverse light/dark cycle (1500 lights on; 0300 lights off) with ad libitum access to food and water. After arrival, rats were given at least 7 days to acclimate to the animal facility prior to surgery. All protocols and animal procedures were conducted in accordance with National Institutes of Health Guide for the Care and Use of Laboratory Animals under supervision of the Institutional Animal Care and Use Committee at Drexel University College of Medicine.

### Drugs

Cocaine hydrochloride was provided by the National Institute on Drug Abuse Drug Supply Program (Research Triangle Park, NC, USA). For self-administration experiments, cocaine hydrochloride was dissolved in 0.9% physiological saline. For FSCV experiments, it was dissolved in artificial cerebrospinal fluid (aCSF).

### Intravenous catheter surgery

Rats were anesthetized using 2.5% isoflurane and implanted with a silastic catheter (ID, 0.012 in OD, 0.025 in. Access Technologies, Skokie, IL) into the right jugular vein for intravenous delivery of cocaine. The catheter was connected to a cannula which exited through the skin on the dorsal surface in the region of the scapulae. Ketoprofen (Patterson Veterinary, Devens, MA; 5mg/kg s.c.) and Enrofloxacin (Norbrook, Northern Ireland; 5 mg/kg s.c.) were provided at the time of surgery and a second dose was given 12 h later. In addition, antibiotic/analgesic powder (Zoetis, Kalamazoo, MI) was applied around the chest and back incisions. Rats were subsequently singly housed and allowed to recover for 5 days prior to self-administration training. Intravenous catheters were manually flushed with gentamicin (5 mg/kg i.v.; Vedco, St. Joseph, MO) in heparinized saline every day during recovery to maintain catheter patency and prevent infection.

### Self-Administration

Self-administration chambers contained two levers – one designated active and the other as inactive. All measures were recorded using custom created Ghost Software (Bernosky-Smith et al., 2016). Each rat underwent one cocaine self-administration session per day from 1000-1600.

### Acquisition

Rats were first trained to self-administer cocaine for 6 h/day under a fixed ratio 1 (FR1) schedule whereby a single active lever response initiated an intravenous injection of cocaine (0.75 mg/kg, infused over 2.5 s - 4.5 s) paired with a cue light above the active lever. Each active lever press was followed by a 20-sec timeout during which the levers retracted. Response on the inactive lever were recorded but otherwise had no consequence. The cocaine dose was chosen based on previous cocaine self-administration studies (Zimmer et al., 2012; Calipari et al., 2013; Alonso et al., 2022). Acquisition criteria for moving onto IntA were ≥ 40 infusions in two consecutive sessions within 10 days of beginning self-administration training.

### Intermittent Access (IntA)

Following acquisition, rats were switched to the IntA schedule of reinforcement (Zimmer et al., 2012). Rats on the IntA schedule had access to cocaine for 5 min followed by a 25-min timeout during which both levers retracted. This 30-min trial repeated for 12 trials per session for a total of 6 h per day. Active lever responses resulted in a single intravenous injection of cocaine (0.375 mg/kg), paired with a cue light above the active lever. Inactive lever responses were recorded but had no consequence. Unlike training sessions, there was no timeout following active lever presses during the 5-min access period on the IntA schedule (Alonso et al., 2022). Rats underwent IntA for 7 consecutive days before moving to abstinence.

### Abstinence and cue-induced seeking tests

Following the final IntA self-administration session, rats underwent a forced abstinence period of 7 days. During this phase, rats remained in their home cage (except when performing cue-induced drug seeking tests). To assess cocaine seeking, rats performed a cue-induced drug seeking test on AD1 and AD8. Cue-induced seeking tests were 1 h in length and all cues (lights, lever presentation, etc.) were identical to the training sessions, except that active lever responses did not result in a cocaine infusion. Under these conditions, the number of active lever presses are interpreted as a measure of cocaine seeking.

### Ex vivo fast scan cyclic voltammetry (FSCV)

18 h after the AD8 seeking test, rats were anesthetized with 2.5% isoflurane for 5 min and brains were rapidly dissected and transferred to ice-cold, aCSF containing NaCl (126 mM), KCl (2.5 mM), NaH2PO4 (1.2 mM), CaCl2 (2.4 mM), MgCl2 (1.2 mM), NaHCO3 (25 mM), glucose (11 mM), and L-ascorbic acid (0.4 mM), with pH adjusted to 7.4. A vibratome was used to produce 400 µm-thick coronal sections containing the NAc. Slices were transferred to room temperature oxygenated aCSF and left to equilibrate for at least 1 h before being transferred into a recording chamber with aCSF (32°C).

A bipolar stimulating electrode was placed on the surface of the tissue in the NAc, and a carbon fiber microelectrode was implanted between the stimulating electrode leads. Dopamine release was evoked every 3 min using a single electrical pulse (400 µA, 4ms, monophasic) and measured using Demon Voltammetry and Analysis Software (Yorgason et al., 2011). Once baseline dopamine release was stable (3 successive stimulations within <10% variation), the slice was exposed to increasing cocaine concentrations (in µM; 0.3, 1.0, 3.0, 10, 30). FSCV data were analyzed in Demon Voltammetry and Analysis Software using Michaelis-Menten kinetic methods to calculate the maximal rate of dopamine uptake (Vmax) and DAT sensitivity to cocaine (i.e., cocaine-induced inhibition of dopamine uptake; apparent Km) (Yorgason et al., 2011).

### Western Blotting

Synaptosomes were prepared, and membrane fractionation was preformed using a modification of published procedures (Brodnik et al., 2020a; Brodnik et al., 2020b; Alonso et al., 2021). Rats were decapitated and the ventral striatum was dissected and stored at -80C until preparation. Tissue was homogenized in ice-cold lysis buffer (1000ml, 50 mM Tris-HCl, pH 7.4, 1 mM EDTA, 320 mM sucrose) with 1x protease inhibitor cocktail, 1x phosphatase inhibitor cocktail, and 1 mM PMSF. The homogenate was centrifuged at 1,000x g for 5 min at 4°C. The resulting supernatant was recentrifuged at 10,000x g for 20 min at 4°C. The resulting synaptosomal pellet was resuspended with 300ml lysis buffer for Western blot studies. Immunoblotting was performed with rabbit anti-DAT polyclonal antibody (1:1000, EMD Millipore), rabbit anti phospho-DAT polyclonal antibody (1:1000, PhosphoSolutions), and peroxidase-conjugated goat anti-rabbit IgG (H1 L) (1:5000, Jackson ImmunoResearch Laboratories). GAPDH was used as membrane protein control and was determined with rabbit anti-GAPDH polyclonal antibody (1:5000, Thermo Fisher Scientific). Total DAT (tDAT), phosphorylated DAT at threonine 53 (pDAT), and GAPDH immunoblots were quantified by densitometry with ImageQuant LAS4000 (GE Healthcare Bio-Sciences). Data were analyzed and presented as a ratio of each protein of interest to GAPDH, as previously reported (Brodnik et al., 2020a; Brodnik et al., 2020b; Alonso et al., 2021).

## Results

### Intermittent access to cocaine promotes incubation of cocaine seeking in a subset of rats

To examine to what extent IntA to cocaine elicits incubation of cocaine seeking after a short period of abstinence, rats self-administered for 7 consecutive days, followed by cue-induced drug seeking tests on AD1 and AD8 (**Figure 1A**). On average, rats acquired self-administration (2 days of >40 presses) in 2.59 ± 0.37 days (**Figure 1B**) and displayed robust cocaine intake during the 7 days of IntA (**Figure 1C**). A paired Student’s t-test revealed that as a group rats pressed significantly more on AD8 compared to AD1 (*t*(16) = 2.425, *p* < 0.028), indicating robust incubation of seeking (**Figure 1D**). However, only a subset of rats displayed greater pressing on AD8 than on AD1, indicating that not all rats displayed incubation of cocaine seeking. A one-way ANOVA with incubation group (non-incubated vs incubated) as the between-subjects variable and abstinence day (AD1 vs AD8) as the within-subjects variable revealed a significant effect of day (*F*(1,15) = 4.725, *p* < 0.5), a significant day by incubation group interaction (*F*(1,15) = 23.76, *p* < 0.005), but no effect of incubation group (*F*(1,15) = 1.195, *p <* 0.29) on the number of lever presses during cue-induced cocaine seeking tests. Holm-Bonferroni post-hoc tests indicated that incubated rats displayed greater pressing on AD8 compared to AD1, and compared to rats that did not incubate on AD8 (**Figure 1E**).

To assess whether the presence/absence of incubation was a consequence of potential differences in cocaine intake, we compared the number of days to acquire self-administration and patterns of cocaine intake between incubated and non-incubated subgroups. A Student’s t-test showed that there were no differences in the number of days to acquire self-administration between non-incubated vs incubated groups (*t*(15) = 0.149, *p* < 0.88; **Figure 1F**). Further, a two-way mixed design ANOVA with incubation group (non-incubated vs incubated) as the between-subjects variable and day of self-administration (days 1-9) as the within-subjects variable revealed no effect of day (*F*(3.366,50.50) = 0.818, *p* < 0.501) or incubation group (*F*(1,15) = 0.253, *p* < 0.622) and no day x incubation group interaction (F(8,120) = 0.339, *p <* 0.949) on the number of lever presses taken over the course of self-administration (**Figure 1G**). These results suggest that neither acquisition nor intake differed between incubated and non-incubated rats.

**Figure 1.**
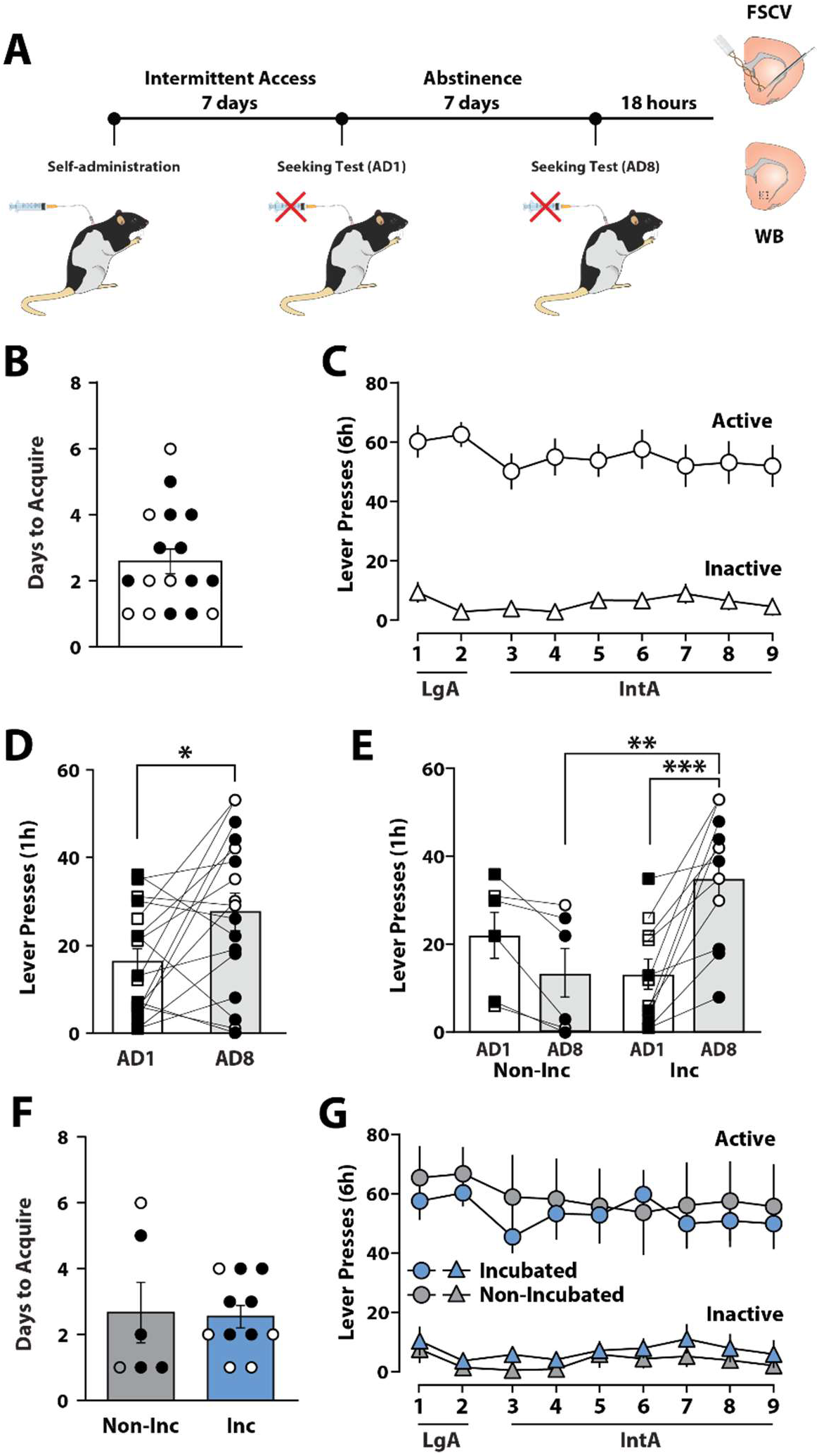
Abstinence from IntA to cocaine promotes incubation of cocaine seeking. **(A)** Experimental timeline for behavioral, fast-scan cyclic voltammetry (FSCV), and Western Blot (WB) data. **(B)** Number of days to acquire cocaine self-administration for all rats. **(C)** Active and inactive lever presses across self-administration sessions for all rats. **(D)** Active lever presses during cue-induced seeking tests for all rats. **(E)** Active lever presses during cue-induced seeking tests for non-incubated (Non-Inc) and incubated (Inc) rats. **(F)** Number of days to acquire cocaine self-administration for Non-Inc and Inc rats. **(G)** Active and inactive lever presses across self-administration sessions for Non-Inc and Inc rats. Data shown as mean±SEM. ○females, ●males. Student’s t-test and two-way ANOVA, \**p* < 0.05, \*\**p* < 0.01, \*\*\**p* < 0.01.

### Incubation of cocaine seeking promotes enhanced dopamine transporter function

To assess if abstinence from IntA to cocaine engendered changes in dopamine transmission, we investigated NAc dopamine release and uptake dynamics 18 h following seeking tests on AD8 (**Figure 1A**). Student’s t-tests indicated that IntA to cocaine resulted in significant increases in dopamine release (t=2.084, *p* < 0.05) and dopamine uptake (t=2.569, *p* < 0.05) compared to cocaine-naive control rats (**Figure 2A-C**). We then asked if these changes in dopamine transmission were specific to rats that developed incubation of cocaine seeking. A one-way ANOVA with incubation group (naive, non-incubated vs incubated) as the between-subjects variable revealed no significant effect of incubation group (*F*(2,26) = 2.365, p > 0.05) on dopamine release, but there was a significant effect of group for dopamine uptake (*F*(2,26) = 4.228, p < 0.05). Holm-Bonferroni post-hoc analyses indicated that incubated rats showed significantly higher dopamine uptake than either naive or non-incubated rats (**Figure 2A,D,E**). These findings indicate that enhanced dopamine uptake during abstinence from IntA to cocaine are specific to rats that develop incubation of seeking.

To assess if IntA and abstinence are associated with changes in dopamine responses to cocaine, we conducted FSCV with bath application of cocaine. A two-way ANOVA with cocaine history (naive vs IntA) as the between subject variable and cocaine concentration (0.3, 1.0, 3.0, 10, and 30 µM) as the within-subject variable showed a significant effect of concentration (Greenhouse-Geisser correction; *F*(2.263, 61.10) = 85.06 p < 0.0001) and a significant cocaine history x concentration interaction (*F*(4, 108) = 4.352, p < 0.005), but no significant effect of cocaine history (*F*(1,27) = 0.1358, p > 0.05) on cocaine-induced changes in dopamine release (**Figure 2F**). Holm-Bonferroni post-hoc tests did not show significant differences at any specific cocaine concentration.

We then tested if IntA and abstinence altered DAT sensitivity to cocaine. A two-way ANOVA with cocaine history (naive vs IntA) as the between-subjects variable and cocaine concentration (0.3, 1.0, 3.0, 10, and 30 µM) as the within-subjects variable revealed a significant effect of cocaine history (*F*(1, 27) = 12.96, p < 0.005), concentration (Greenhouse-Geisser correction; *F* (1.131, 30.53) = 332.8, p < 0.0001), and cocaine history x concentration interaction (*F*(4, 108) = 10.84, p < 0.0001) on DAT sensitivity to cocaine. Holm-Bonferroni post-hoc tests revealed that IntA and abstinence promoted greater DAT sensitivity to cocaine at all but the 10 μM cocaine concentration (**Figure 2G**).

To examine if changes in dopamine responses to cocaine were specific to incubation of cocaine seeking, we again separated cocaine-experienced rats into non-incubated and incubated subgroups. A two-way ANOVA with incubation group (naive, non-incubated, vs incubated) as the between-subject variable and cocaine concentration as the within-subject variable revealed a significant effect of concentration (Greenhouse-Geisser correction; *F*(2.247, 58.43) = 82.20, p < 0.0001) and a significant incubation group x concentration interaction (*F*(8, 104) = 2.580, p < 0.05), but no significant effect of incubation group (*F*(2, 26) = 0.1690, p > 0.05) on cocaine-induced dopamine release (**Figure 2H**). Holm-Bonferroni tests did not show significant differences at any specific cocaine concentration.

Finally, we examined whether DAT sensitivity to cocaine differed between incubated and non-incubated rats. A one-way ANOVA with incubation group (naive, non-incubated, vs incubated) as the between subjects variable and cocaine concentration as the within subjects variable revealed a significant effect of incubation group (*F*(2, 26) = 8.716, p < 0.005), concentration (*F*(1.145, 29.76) = 370.5, p < 0.0001), and a significant incubation group x concentration interaction (*F*(8, 104) = 7.636, p < 0.0001) on DAT sensitivity to cocaine (**Figure 2I**). Holm-Bonferroni tests showed that incubated rats displayed greater DAT sensitivity to cocaine compared to naive rats at all cocaine concentrations and that non-incubated rats showed a significant increase compared to naive rats only at the 1μM cocaine concentration.

**Figure 2.**
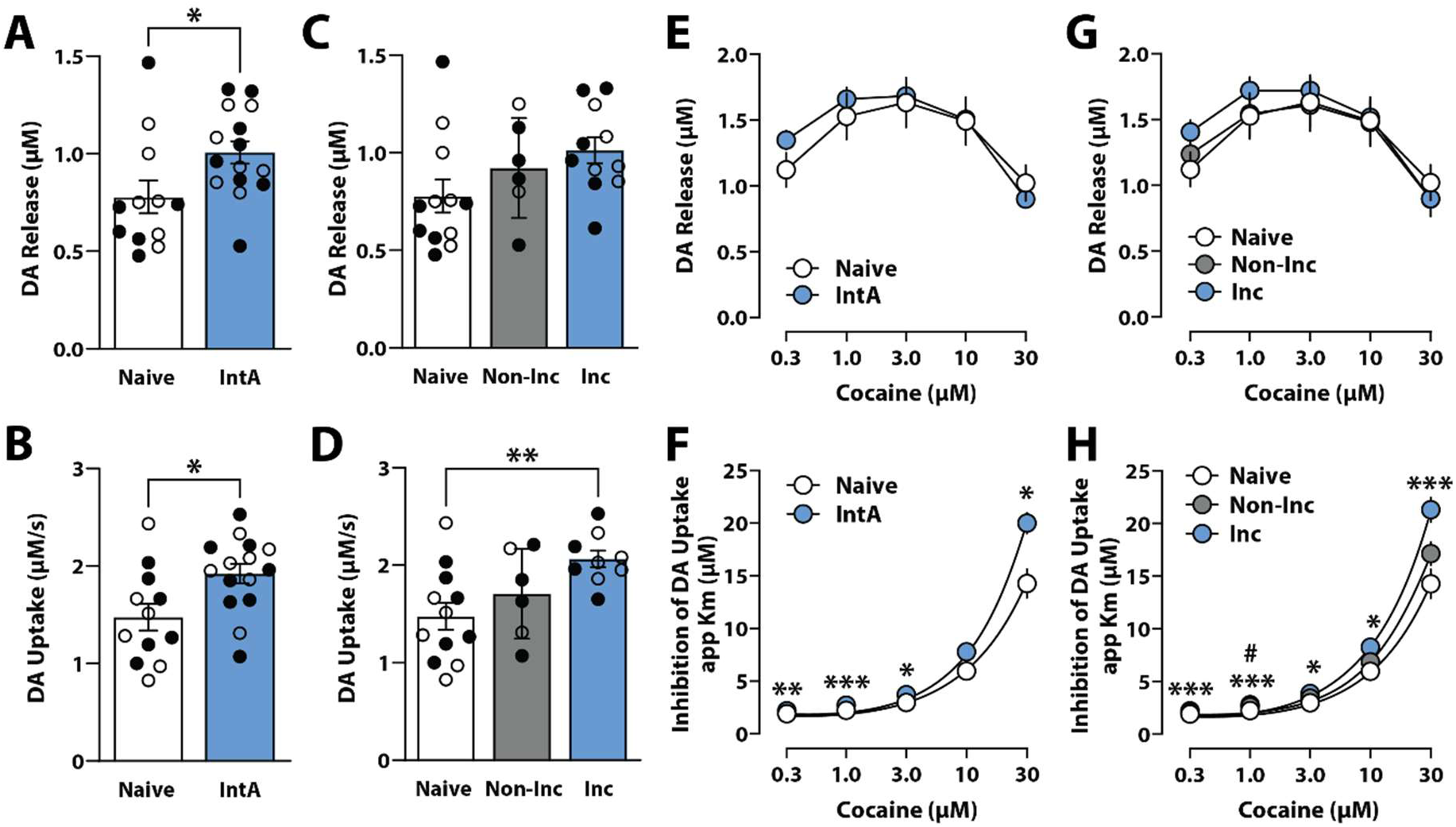
Abstinence from IntA to cocaine enhances dopamine release and uptake and DAT sensitivity to cocaine in the nucleus accumbens. **(A)** Baseline dopamine release and **(B)** dopamine uptake for naive and all intermittent access (IntA) rats. **(C)** Baseline dopamine release and **(D)** dopamine uptake for naive, non-incubated (Non-Inc), and incubated (Inc) rats. **(E)** Cocaine-induced dopamine release and **(F)** Inhibition of dopamine uptake (i.e, DAT sensitivity to cocaine; app Km) for naive and all IntA rats. **(G)** Cocaine-induced dopamine release and **(H)** inhibition of dopamine uptake for naive, Non-Inc, and Inc rats. Data shown as mean±SEM. ○females, ●males. Student’s t-test, one-way ANOVA, or two-way ANOVA, \**p* < 0.05, \*\**p* < 0.01, \*\*\**p* < 0.001, IntA vs naive or Inc vs Naive. ^#^*p* < 0.05, Non-Inc vs Naive.

### Incubation of cocaine seeking promotes enhanced DAT expression and phosphorylation

We next asked if observed enhancements in DAT function were associated with changes in DAT surface expression or phosphorylation. NAc tissue from naive controls and rats that underwent IntA to cocaine was analyzed with Western blotting for total membrane DAT (tDAT) and DAT phosphorylated at Challasivakanaka S (pDAT; **Figure 1A**). Student’s t-tests revealed modest, non-significant increases in total DAT expression (t(16) = 1.202 p > 0.05) and pDAT (t(16) = 0.873 p > 0.05) between naive and IntA rats (**Figure 3A-C**). Given our earlier finding that dopamine dynamics differed between incubated and non-incubated rats, we examined tDAT and pDAT separately in non-incubated and incubated subgroups. A one-way ANOVA with incubation group (naive, non-incubated vs incubated) as the between subjects variable revealed a significant effect of incubation group on tDAT (*F*(2, 15) = 5.36, p < 0.05) and pDAT (*F*(2,15) = 4.151, p < 0.05) expression (**Figure 3A,D,E**). Holm-Bonferroni tests further demonstrated that incubated rats exhibited significantly greater tDAT and pDAT levels than naive controls.

**Figure 3.**
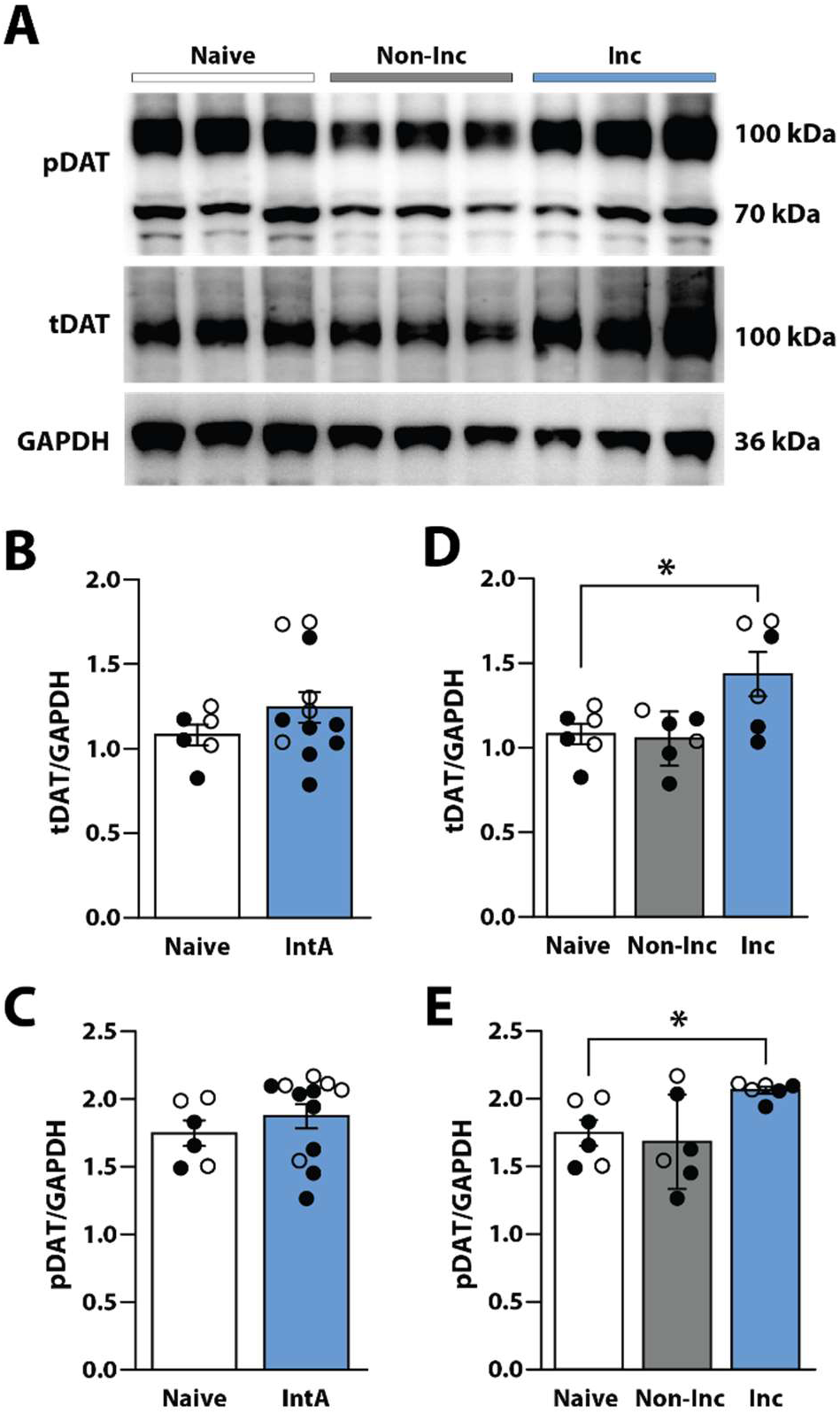
Incubated rats display reduced DAT and pDAT expression. **(A)** Example Western blots. **(B)** Quantification of total membrane DAT (tDAT), and **(C)** phosphorylated DAT (pDAT) over GAPDH for naive and intermittent access (IntA) rats. **(D)** Quantification of tDAT and **(E)** pDAT over GAPDH for naive, non-incubated (Non-Inc) and incubated (Inc) rats. Data shown as mean±SEM. ○females, ●males. Student’s t-test, \**p* < 0.05.

### Hcrtr1 blockade early in abstinence prevents incubation of cocaine seeking

Extensive evidence indicates that disrupting Hcrtr1 signaling reduces behavioral and dopamine responses to cocaine (España et al., 2010; Bentzley and Aston-Jones, 2015; Prince et al., 2015; Bernstein et al., 2017; Levy et al., 2017; Shaw et al., 2017; Black et al., 2023). Consequently, we investigated the effects of Hcrtr1 blockade on incubation of cocaine seeking and dopamine adaptations following abstinence from IntA to cocaine. Separate groups of female and male rats underwent IntA self-administration and 7 days of abstinence (**Figure 4A**). Immediately, after the seeking test on AD1, rats received a single intraperitoneal (i.p.) injection of vehicle or (20 mg/kg) of the Hcrtr1 antagonist RTIOX-276 (Perrey et al., 2013). A Student’s t test revealed no differences in days to acquire cocaine self-administration between rats that would eventually receive vehicle or RTIOX-276 (t(16) = 0.618, p < 0.545; **Figure 4B**). Similarly, a two-way ANOVA with future treatment (vehicle vs RTIOX-276) as the between subjects variable and day of self-administration (days 1-9) as the within-subjects variable revealed no effect of future treatment (*F*(1, 16) = 0.7726, p < 0.392), day (Greenhouse-Geisser Correction; *F*(3.378, 54.05) = 1.706, p < 0.171), or a future treatment x day interaction (*F*(8, 128) = 1.546, p < 0.147) on the number of injections taken over the course of self-administration (**Figure 4C**).

We then assessed whether HcrtR1 blockade affected incubation of cocaine seeking. A two-way ANOVA with treatment (vehicle or RTIOX-276) as the between-subjects variable and seeking test day (AD1 vs AD8) as the within-subjects variable revealed no effect of treatment (*F*(1, 16) = 1.312, p < 0.269), but did reveal an effect of day (*F*(1, 16) = 8.454, p < 0.01) and a treatment x day interaction (*F*(1, 16) = 6.582, p < 0.05) on active lever presses. Holm-Bonferroni tests indicated a significant increase in active lever presses on AD8 compared to AD1 in rats treated with vehicle, indicating the expected incubation of cocaine seeking observed following IntA to cocaine. Rats treated with RTIOX-276, however, did not show increases in cocaine seeking between AD8 and AD1 and displayed significantly lower pressing on AD8 than vehicle-treated rats. These results indicate that RTIOX-276 prevented incubation of cocaine seeking (**Figure 4D**).

**Figure 4.**
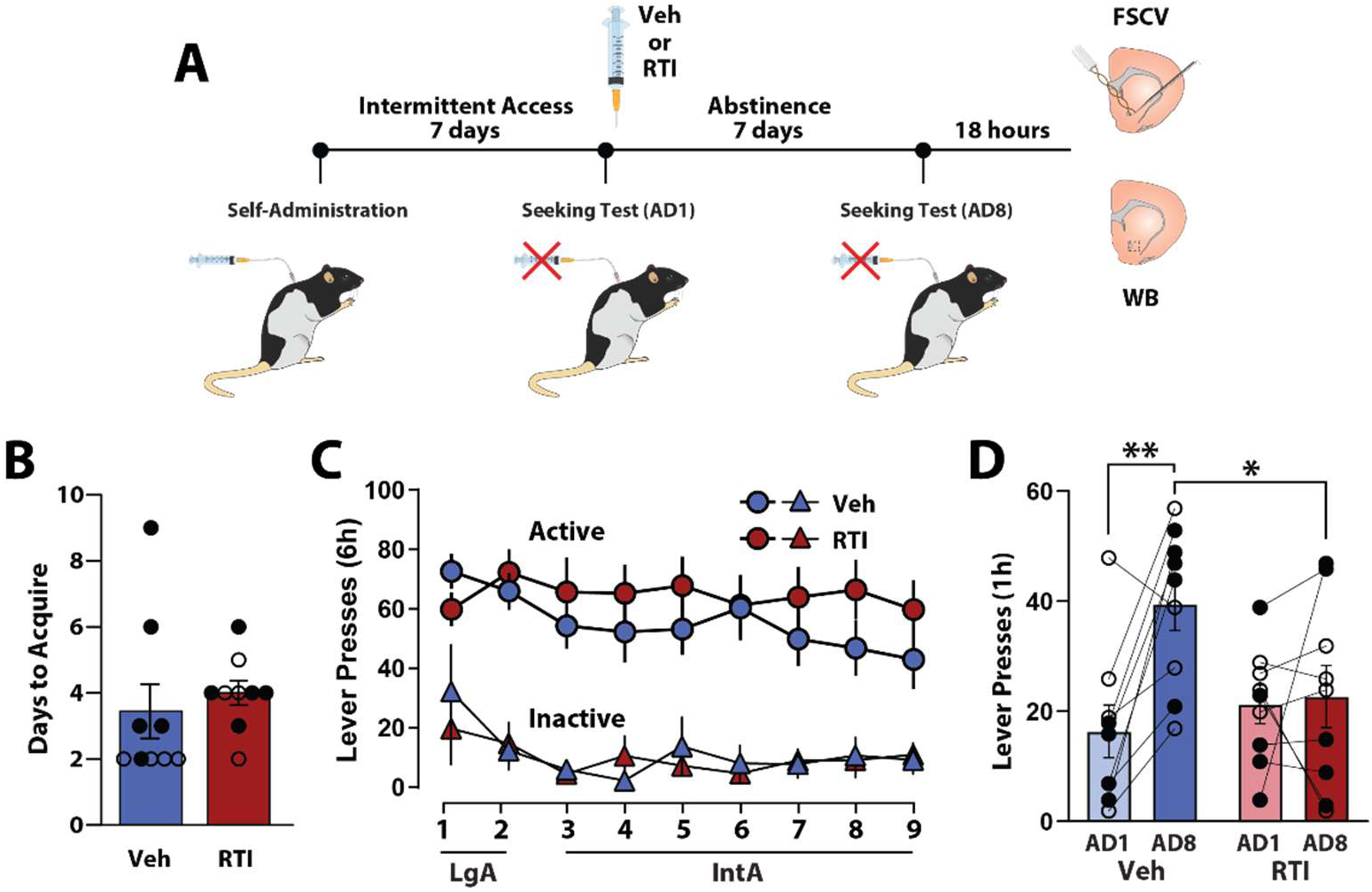
HCRTr1 blockade prevents incubation of cocaine seeking. **(A)** Experimental timeline for behavioral, fast-scan cyclic voltammetry (FSCV), and Western Blot (WB) data. **(B)** Number of days to acquire cocaine self-administration in rats treated with vehicle (Veh) or 20mg/kg RTIOX-276 (RTI). **(C)** Active and inactive lever presses during self-administration. **(D)** Active lever presses during cue-induced seeking tests. Data shown as mean±SEM. ○females, ●males. Two-way ANOVA, \**p* < 0.05. \*\**p* < 0.01.

### Hcrtr1 blockade normalizes dopamine transmission and DAT biochemistry

To determine if reductions in cocaine seeking following Hcrtr1 antagonism were associated with normalization of dopamine transmission, we measured dopamine release and uptake in the NAc 18 h following seeking tests on AD8 (**Figure 4A**). A Student’s t-test revealed that Hcrtr1 antagonism after cue-induced seeking tests on AD1 significantly reduced dopamine release (t(15)= 2.744, *p* < 0.05) and dopamine uptake (t(15) = 2.841, *p* < 0.05) compared to vehicle treatment (**Figure 5A and B**).

To assess whether antagonism of Hcrtr1 reduced dopamine responses to cocaine, we conducted FSCV with bath application of cocaine. A two-way mixed design ANOVA with treatment (vehicle vs RTIOX-276) as the between-subjects variable and cocaine concentration (0.3, 1.0, 3.0, 10, 30 μM) as the within-subjects variable showed no significant effect of treatment (*F*(1, 15) = 2.810, *p* < 0.05), but there was a significant effect of concentration (Greenhouse-Geisser correction; *F*(2.130, 31.95) = 56.90, *p* < 0.0001) and a significant treatment x concentration interaction (Greenhouse-Geisser correction; *F*(4, 60) = 3.487, *p* < 0.05) on cocaine-induced dopamine release (**Figure 5D**). Holm-Bonferroni tests indicated that RTIOX-276 significantly reduced dopamine release only at the 0.3 μM concentration.

Next, we examined DAT sensitivity to cocaine using a two-way ANOVA with treatment (vehicle vs RTIOX-276) as the between-subjects variable and cocaine concentration (0.3, 1.0, 3.0, 10, and 30 µM) as the within-subjects variable. This analysis revealed a significant effect of treatment (*F*(1, 15) = 9.918, *p* < 0.01), concentration (Greenhouse-Geisser correction; *F*(1.055, 15.82) = 135.5, *p* < 0.001), and treatment x concentration interaction (*F*(4, 60) = 9.499, *p* < 0.0001) on DAT sensitivity to cocaine (**Figure 5E**). Holm-Bonferroni tests revealed that Hcrtr1 antagonism significantly reduced DAT sensitivity to cocaine at the 10 and 30 µM cocaine concentrations.

Lastly, we examined RTIOX-276 prevented changes in DAT expression observed following incubation of cocaine seeking. Student’s t-test showed that RTIOX-276 significantly reduced both tDAT (t(11) = 2.892, *p* < 0.05) and pDAT (t(11) = 2.841, *p* < 0.05) expression (**Figure 5F-G**), further suggesting that changes in dopamine transmission are linked to changes in phosphorylation state of the DAT.

**Figure 5.**
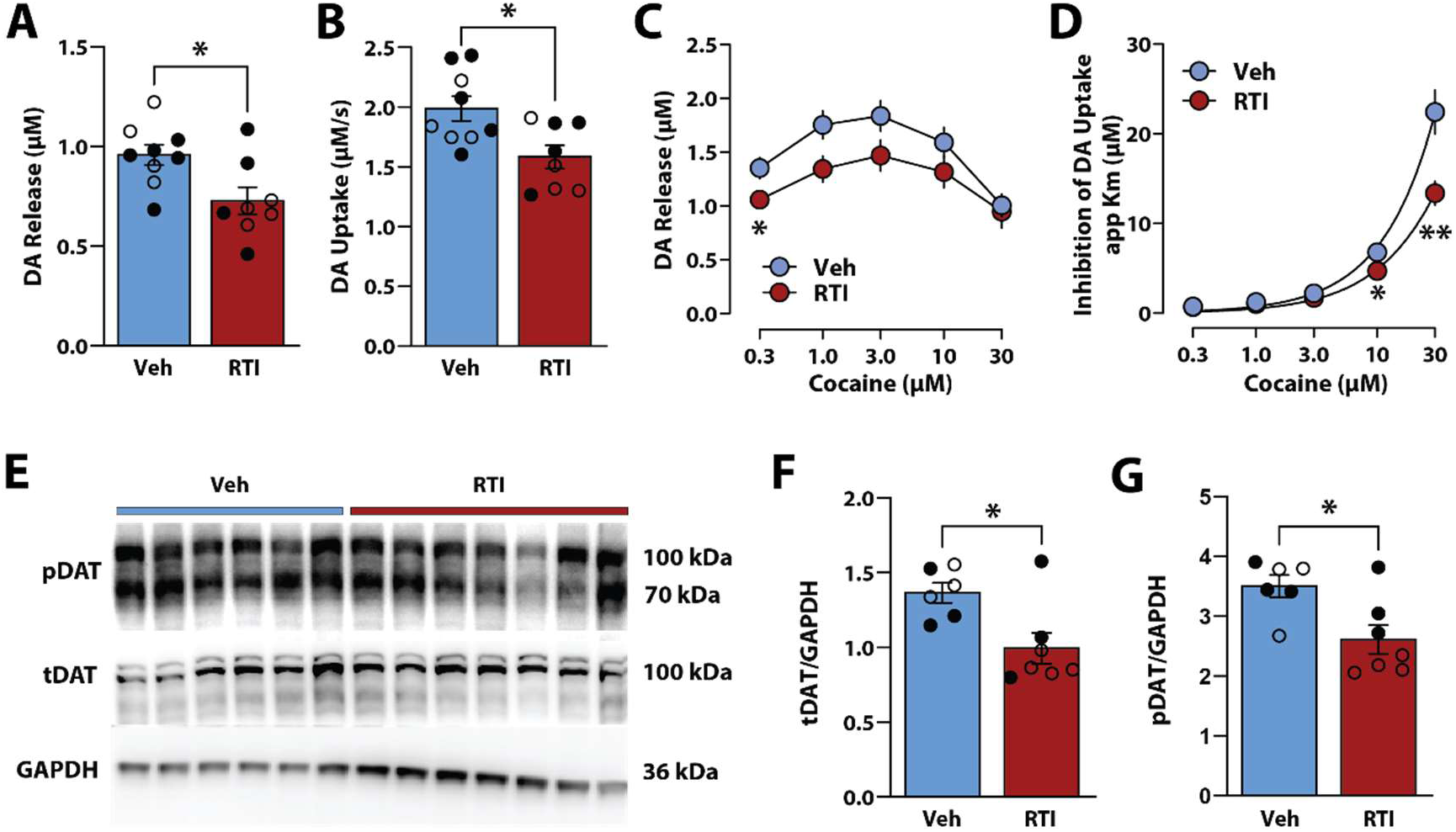
HcrtR1 blockade normalizes dopamine transmission and DAT expression. **(A)** Baseline dopamine release and **(B)** dopamine uptake for rats treated with vehicle (Veh) or 20mg/kg RTIOX-276 (RTI). **(C)** Cocaine-induced dopamine release and **(D)** inhibition of dopamine uptake (i.e, DAT sensitivity to cocaine; app Km). **(E)** Example Western blots. **(F)** Quantification of total membrane DAT (tDAT), and **(G)** phosphorylated DAT (pDAT) over GAPDH. Data shown as mean±SEM. ○females, ●males. Student’s t-test or two-way ANOVAs, \**p* < 0.05, \*\**p* < 0.01.

## Discussion

In the present studies we examined whether incubation of cocaine seeking is associated with changes in dopamine transmission and to what extent Hcrtr1 blockade early in abstinence reduces incubation of cocaine seeking and normalized dopamine transmission later in abstinence. Results indicated that IntA to cocaine engenders incubation of cocaine seeking as a group, but that only subset of rats displayed incubation. We also observed that increases in dopamine release and uptake and dopamine responses to cocaine were increased preferentially in rats that showed an incubation of cocaine seeking. Furthermore, we found that a single injection of the Hcrtr1 antagonist RTIOX-276 on the first day of abstinence prevented incubation of cocaine seeking and aberrant dopamine adaptations one week later. Together, these findings suggest that Hcrtr1 antagonism early in abstinence may serve as a therapeutic for reducing cocaine seeking, thus decreasing the likelihood for relapse

### IntA followed by a brief abstinence engenders incubation of cocaine seeking

Incubation of cocaine seeking or the tendency of cue-induced cocaine seeking to increase throughout abstinence has received significant attention as a key factor leading to relapse (Grimm et al., 2001; Grimm et al., 2003; Bienkowski et al., 2004; Lu et al., 2004; Guillem et al., 2014; Calipari et al., 2015; Chen et al., 2015; Parvaz et al., 2016; Gu, 2018; Alonso et al., 2022). In the present study, rats were allowed IntA to cocaine, which promotes elevated cocaine intake rates and results in high peak concentrations of the drug (Zimmer et al., 2012). This exposure is thought to better model the binge-like patterns of cocaine consumption seen in humans (Beveridge et al., 2012; Zimmer et al., 2012; Alonso et al., 2022) and possibly reflects an increase in the perceived scarcity of the drug, thereby potentiating its reward value (Acuff et al., 2023). We previously showed that IntA promotes robust incubation of cocaine seeking after 28 days of abstinence (Alonso et al., 2022). However, given that cocaine-associated adaptations appear early in abstinence (Von Diemen et al., 2014; Kumaresan et al., 2023) and can be predictive of relapse risk (McHugh et al., 2014; Poireau et al., 2022), here we assessed if IntA to cocaine also produces incubation of cocaine seeking earlier in abstinence. Results indicated that as early as 7 days of abstinence, IntA to cocaine promoted robust incubation of seeking. These findings support the potential for IntA to model behavioral changes that occur at early points in abstinence, and thus provides a platform for testing therapeutics that can be administered early in abstinence to curtail the development of incubation of cocaine seeking.

### Changes in NAc dopamine transmission are tied to incubation of cocaine seeking

Exposure to cocaine leads to plasticity in the mesolimbic dopamine pathway, including long-lasting potentiation of excitatory input to VTA dopamine neurons (Bellone and Lüscher, 2006; Chen et al., 2008; Yuan et al., 2013). We and others have also shown functional changes in dopamine transmission in the NAc during abstinence from cocaine (Calipari et al., 2015; Alonso et al., 2022), including increases in dopamine uptake rate and DAT sensitivity to cocaine following 28 days of abstinence (Alonso et al., 2022). Consistent with these findings, here we observed increased dopamine release, dopamine uptake and DAT sensitivity to cocaine after 7 days of abstinence from IntA to cocaine. Despite these observations, it was not clear if these dopamine adaptations were a consequence of IntA cocaine exposure alone, abstinence from IntA to cocaine, or whether they were tied to incubation of cocaine seeking. To address this question, leveraged the individual differences observed in cocaine seeking behavior by separating IntA rats into those that developed incubated seeking and those that did not. This split reflects findings from human studies demonstrating that only a proportion of individuals who use cocaine go on to develop cocaine use disorder (Schwartz et al., 2022). Incubated and non-incubated rat groups did not differ in the amount of cocaine consumed or other measures of cocaine self-administration prior to abstinence, but only the incubated rats displayed increases in dopamine release and uptake and exaggerated DAT sensitivity to cocaine. In fact, non-incubated rats demonstrated similar measures of dopamine transmission as cocaine-naive controls. While chronic cocaine use has been associated with persistent changes in mesolimbic dopamine function in both rodents and humans (Volkow et al., 1996; Ferris et al., 2012; Calipari et al., 2013; Calipari et al., 2014), to our knowledge, this is the first study to demonstrate that dopamine disruptions are preferentially observed in subjects that express incubation of cocaine seeking.

### HCTRr1 blockade prevents incubation of seeking and normalizes dopamine transmission in the NAc

A principle challenge in treating cocaine use disorder is reducing intensification of cocaine craving during periods of abstinence in an effort to decrease the risk of relapse (Grimm et al., 2001; Lu et al., 2004). Given our observation that incubation of cocaine seeking was accompanied by dopamine adaptations, we hypothesized that normalizing these dopamine adaptations would prevent development of incubation of cocaine seeking. Extensive evidence indicates that the Hcrts influence cocaine-associated behavior and that these effects involve actions on dopamine transmission, including changes in dopamine uptake, pDAT, and dopamine responses to cocaine (Borgland et al., 2006; España et al., 2010; Moorman and Aston-Jones, 2010; Smith et al., 2010; Bentzley and Aston-Jones, 2015; Brodnik et al., 2015; Bernstein et al., 2017; Levy et al., 2017; Shaw et al., 2017; Martin-Fardon et al., 2018; Brodnik et al., 2020a; Black et al., 2023). Despite this evidence, the therapeutic potential of HcrtR1 blockade for treating incubation of cocaine seeking has not been previously reported. Consequently, in these studies we administered the Hcrtr1 antagonist, RTIOX-276, on the first day of abstinence – a timepoint chosen to maximize potential therapeutic benefit by curtailing development of incubation of cocaine seeking. Additionally, because adherence to treatments for substance use disorder tends to wane over the course of abstinence an effective treatment for reducing craving early in abstinence would be highly valuable for reducing rates of relapse (Morasco et al., 2011; Stoner et al., 2015). We found that administration of RTIOX-276 immediately following seeking tests on AD1 significantly decreased cue-induced cocaine seeking one week later on AD8. Moreover, when examining if these behavioral effects were linked with changes in dopamine transmission, we observed that RTIOX-276 reduced the expected enhancement in dopamine transmission observed following IntA to cocaine.

While most prior studies on Hcrts and cocaine-associated behavior have been restricted to the acute, onboard pharmacological effects of HcrtR1 antagonists, a few studies have reported lasting effects of HcrtR1 antagonists in the context of drug abuse (Ishii et al., 2005; Winrow et al., 2010; Brodnik et al., 2020a; Mohammadkhani et al., 2020). For example, evidence from our laboratory indicates that HcrtR1 blockade alters cocaine self-administration and mesolimbic dopamine transmission for up to 24 h after treatment (Brodnik et al., 2020a). While the mechanism remains elusive, HcrtR1 blockade may exert lasting effects on dopamine transmission in the NAc through acute alterations in dopamine neuron activity within the VTA. Not only does HcrtR1 blockade VTA dopamine neuron activity (Narita et al., 2006; Muschamp et al., 2007; Moorman and Aston-Jones, 2010; Muschamp et al., 2014), which decreases dopamine responses to cocaine in the NAc (España et al., 2010), but we have also shown that chemogenetic inhibition of VTA dopamine neurons leads to reductions in DAT sensitivity to cocaine and decreases in DAT phosphorylation at threonine 53 in the NAc (Brodnik et al., 2020b). Given that cocaine exposure engenders plastic changes in glutamate receptors of VTA neurons within hours of cocaine exposure (Ungless et al., 2001; Brown et al., 2010) and that dopamine release is also significantly increased shortly after cocaine intake ceases (Marinelli et al., 2003), it is possible that Hcrtr1 antagonism might prevent or reverse this dopamine plasticity during a sensitive period early in abstinence by attenuating increases in dopamine firing. Moreover, given the prominent actions of Hcrts in signaling arousal states (Tyree et al., 2018), it is also possible that Hcrtr1 blockade following the first cue-induced seeking test disrupts cortical or limbic integration of the seeking test context and strength of cocaine-cue pairings (Capriles et al., 2003; Perry et al., 2014; Hill-Bowen et al., 2021; Acuff et al., 2023), which would be expected to diminish the salience of cocaine cues.

### Alterations in DAT expression and phosphorylation may explain changes in dopamine transmission

Accumulating evidence suggests that phosphorylation of DAT at threonine 53 is associated with faster dopamine uptake, increased DAT sensitivity to cocaine, and increased motivation for cocaine (Foster et al., 2012; Calipari et al., 2017; Challasivakanaka et al., 2017; Foster and Vaughan, 2017; Brodnik et al., 2020a). Here we observed higher pDAT levels and greater total membrane DAT expression preferentially in rats that displayed incubation of cocaine seeking. These effects matched our FSCV findings in which only rats that displayed incubation of cocaine seeking displayed increases in dopamine uptake rate and DAT sensitivity to cocaine. The effects of HcrttR1 blockade on dopamine transmission are also consistent with changes in DAT biochemistry, with RTIOX-276 reducing both total membrane DAT expression and pDAT after abstinence from IntA to cocaine. When considered together, these findings suggest that changes in dopamine uptake and DAT sensitivity to cocaine observed in rats that incubated their seeking may be mediated by increased DAT expression and phosphorylation, similar to what we have suggested previously (Brodnik et al., 2020a; Brodnik et al., 2020b; Alonso et al., 2021). The observed results also support the premise that RTIOX-276 reduces cocaine seeking by preventing these dopamine adaptations.

Lastly, in addition to DAT phosphorylation state or total membrane expression, other mechanisms may contribute to alterations in DAT function. For example, several reports suggest that shifts in the proportion of inward/outward facing DATs (Liang et al., 2009), changes in oligomer/monomer ratios (Zhen et al., 2015; Siciliano et al., 2018), dimerization with sigma receptor (Hong et al., 2017), or phosphorylation of DAT serine sites (Khoshbouei et al., 2004; Moritz et al., 2013) can impact the efficiency of dopamine uptake. Therefore, although our findings demonstrate a link between DAT function, DAT expression, and DAT phosphorylation at threonine 53, other mechanisms are likely to be involved.

## Summary

In the present studies, we demonstrated that IntA to cocaine engenders dopamine adaptations that underlie incubation of cocaine seeking and that HcrtR1 antagonism early in abstinence prevents dopamine adaptations and exerts lasting reductions in cue-induced cocaine seeking. Given that early abstinence represents a particularly vulnerable period for treating cocaine use disorder, these findings suggest that Hcrtr1 antagonism early in abstinence – when cocaine craving begins to develop – may serve as a potentially valuable pharmacotherapeutic option for decreasing craving and reducing the propensity for relapse.

## Acknowledgements

We thank the NIDA drug supply program for donating the cocaine hydrochloride. This work was supported by the National Institute on Drug Abuse grant R01DA039100 (RAE).

## Author Contributions

PJC, conceptualized and designed the studies, conducted the self-administration and FSCV data collection, analyzed and interpreted the data, wrote the original draft, and edited the content. VMM, conducted the self-administration and FSCV data collection, wrote and edited the content. SD, performed western blotting and analyzed the data. SK, conceptualized and designed the studies, analyzed and interpreted the data, wrote and edited the content. RAE, conceptualized and designed the studies, analyzed and interpreted the data, edited the original draft, and edited the content.

